# Considering variance in pollinator responses to stressors can reveal potential for resilience

**DOI:** 10.1101/2021.12.24.474118

**Authors:** Amélie Cabirol, Tamara Gómez-Moracho, Coline Monchanin, Cristian Pasquaretta, Mathieu Lihoreau

## Abstract

1. Environmental stressors have sublethal consequences on animals, often affecting the mean of phenotypic traits in a population. However, potential effects on variance are poorly understood. Since phenotypic variance is the basis for adaptation, any influence of stressors may have important implications for population resilience.
2. Here we explored this possibility in insect pollinators by analysing raw datasets from 24 studies (6,913 bees) in which individuals were first exposed to stressors and then tested for cognitive tasks.
3. While all types of stressors decreased the mean cognitive performance of bees, their effect on variance was complex. Focusing on 15 pesticide studies, we found that the dose and the mode of exposure to stressors were critical. At low pesticide doses, cognitive variance decreased following chronic exposures but not for acute exposures. Acute exposure to low doses thus seems less damaging at the population level. In all cases however, the variance decreased with increasing doses.
4. *Policy implications*. Current guidelines for the authorization of plant protection products on the European market prioritize acute over chronic toxicity assessments on non-target organisms. By overlooking the consequences of a chronic exposure, regulatory authorities may register new products that are harmful to bee populations. Our findings thus call for more research on stress-induced phenotypic variance and its incorporation to policy guidelines to help identify levels and modes of exposure animals can cope with.

## Introduction

Human activities have led to a dramatic increase in the extinction rates of animal species (Barnosky *et al*., 2011; Dirzo *et al*., 2014; Wagner, 2020). Associated stressors have partly been identified and act synergistically (Brook, Sodhi and Bradshaw, 2008; Dirzo *et al*., 2014; Sánchez-Bayo and Wyckhuys, 2019; Siviter *et al*., 2021). These include habitat loss, pollutions, and the introduction of invasive species. These factors add up to the ones naturally encountered by animals in their environment, such as the presence of predators, pathogens, and parasites. Given the raising number of species threatened with extinction (Barnosky *et al*., 2011; Sánchez-Bayo and Wyckhuys, 2019), it has become urgent to understand how animal populations can cope with human-induced stressors in order to orient policies towards an efficient regulation of activities affecting the biodiversity.

Many of these stressors do not kill animals, but nevertheless significantly impact their fitness through inaccurate behaviour or reduced reproduction (Klein et al. 2017). Measuring these sublethal effects of stressors on populations is difficult because of the technical challenge of monitoring large numbers of animals and tease apart the many confounding factors linked to field conditions. Most studies have thus focused on the effects of stressors on individual animals using controlled laboratory setups to measure single phenotypic traits, such as cognition or reproduction (Badyaev 2005). Yet, the relevance of such risk assessment methods compared to field population-level studies has been questioned as mismatching conclusions often emerged from the two approaches (Thompson and Maus, 2007; Henry *et al*., 2015). Even though stressors may affect individual phenotypic traits in the lab, life in a natural, sensory, and socially enriched environment can buffer or amplify these effects (Wright and Conrad, 2008; Henry *et al*., 2015; Lambert *et al*., 2016; Cabirol *et al*., 2017).

Studies investigating the impact of stressors on phenotypic traits often report shifts in their means at the population level. Agrochemicals, for instance, were shown to reduce food consumption and delay migration in songbirds (Eng, Stutchbury and Morrissey, 2019), to alter endocrine functions in amphibians and fish (Mann *et al*., 2009; Besson *et al*., 2020), and to reduce learning performance in bees (Siviter *et al*., 2018). We therefore argue that studying how stressors affect the variance of these traits will provide important complementary information about the severity of stressors on animal populations and may reconcile results obtained in the lab and in the field.

It is well recognized that animals exhibit variability in behavioural and physiological responses to stressors (Ebner and Singewald, 2017; Mazza *et al*., 2019). Some individuals may better cope with particular stressors than others. Thus understanding how this variance in stress-response is affected at the population level is crucial for risk assessment (Nakagawa et al. 2015). If the variance is low in the population following stressor exposure, all individuals may suffer the consequences associated with the altered phenotype. On the contrary, if the variance remains high in the population, even though the mean is affected, some individuals may still exhibit an adaptive phenotype. In some cases, stressors may even increase phenotypic variance, a phenomenon suggested to promote the evolutionary diversification of species (Badyaev, 2005). Stressors act as agents of selection and stress-induced variation should therefore be considered when assessing the resilience of a population to a particular stressor (Hoffmann and Merilä, 1999).

Here we highlight the importance of studying the phenotypic variance in animal populations exposed to stressors. To support this claim, we analysed the effect of stressors on the mean and variance of cognitive performances in bees. We focused on honey bees (*Apis*) and bumblebees (*Bombus*), as they are arguably the most studied pollinators. They are also known to be affected by multiple natural and human-induced stressors, and in particular pesticides (Potts *et al*., 2010; Goulson *et al*., 2015). Honey bees and bumblebees live in colonies with a division of labour and are therefore characterized by an important level of inter-individual behavioural and cognitive variability (Jeanson and Weidenmüller, 2014). Foragers, in particular, have evolved a rich cognitive repertoire enabling them to locate and recognise plant resources, handle them, and navigate back to their hive to unload food for the colony (Chittka, 2017). One of the most reported sublethal effect of stressors on bees is the decrease in their cognitive performance (learning and memory), which has been associated with a decreased foraging success and colony survival (Klein *et al*., 2017). A recent meta-analysis confirmed that exposure to neonicotinoid pesticides at field-realistic doses, either in acute or chronic exposure, consistently alter the mean olfactory learning and memory performance of bees (Siviter *et al*., 2018). However, the impact of stressor intensity (dose and duration) on the variance of the learning performance was not analysed. We therefore explored these effects by analysing the raw datasets from 24 studies that assessed bee cognition applying olfactory and visual learning protocols in either an appetitive or aversive context. Although a decreased cognitive performance was expected in stressed bees, we predicted that the effect of stressors on the variance would depend on the stressor intensity, which would help estimate the hazardous nature of a given stressor.

## Materials and methods

### Search and selection of datasets

The search for scientific publications falling within the scope of our research question was performed in July 2020 using the PubMed database. The keywords used for the search were (“Stressor” OR “Pesticide” OR “Parasite”) AND (“Cognition” OR “Learning”) AND (“Pollinators” OR “Bees”). A total of 71 studies were found, of which 22 met our inclusion criteria. Two datasets belonging to the authors of this study were also included as they filled the inclusion criteria. These studies measured the impact of stressors on the cognitive performance of bees. A summary of the studies is given a Table 1.

**Table 1:** Summary of the 24 studies used.

| Table 1: Summary of the 24 studies used. |  |  |  |
| --- | --- | --- | --- |
| Stressor | Genus | Exposure type | Reference |
| Pesticide | *Apis* | Acute | (Ludicke and Nieh, 2020) |
| Pesticide | *Apis* | Acute | (Hesselbach and Scheiner, 2018) |
| Pesticide | *Apis* | Acute | (Urlacher *et al.*, 2016) |
| Pesticide | *Apis* | Acute, | (Tan *et al.*, 2015) |
| Pesticide | *Apis* | Chronic | (Mustard *et al.*, 2020) |
| Pesticide | *Apis* | Chronic | (Tan *et al.*, 2017) |
| Pesticide | *Apis, Bombus* | Acute | (Siviter *et al.*, 2019) |
| Pesticide | *Bombus* | Acute | (Muth *et al.*, 2019) |
| Pesticide | *Bombus* | Acute, chronic | (Stanley, Smith and Raine, 2015) |
| Pesticide | *Bombus* | Chronic | (Smith *et al.*, 2020) |
| Pesticide | *Bombus* | Chronic | (Lämsä *et al.*, 2018) |
| Pesticide | *Bombus* | Chronic | (Phelps *et al.*, 2018) |
| Pesticide,<br>coexposure | *Apis* | Chronic | (Colin, Plath, *et al.*, 2020) |
| Parasite | *Bombus* | Acute | Gomez-Moracho *et al.* (2021) |
| Parasite | *Bombus* | Acute | (Martin, Fountain and Brown, 2018) |
| Pollution | *Apis* | Acute | Monchanin *et al.* (unpublished) |
| Pollution | *Apis* | Acute | (Monchanin, Drujont, *et al.*, 2021) |
| Pollution | *Apis* | Acute | (Leonard *et al.*, 2019) |
| Pollution | *Apis* | Chronic | (Monchanin, Blanc-brude, *et al.*,<br>2021) |
| Other | *Apis* | Acute | (Wang *et al.*, 2016) |
| Other | *Apis* | Acute | (Shepherd *et al.*, 2018) |
| Other | *Apis* | Chronic | (Shepherd *et al.*, 2019) |
| Coexposure | *Apis, Bombus* | Acute/Chronic | (Piironen and Goulson, 2016) |
| Coexposure | *Bombus* | Acute/Chronic | (Piironen *et al.*, 2016) |

#### Cognitive tasks

We focused on cognitive data from bees exposed to stressors during their adult life, as bees treated as larvae might be more sensitive to stressors (i.e. pesticides; Siviter et al. 2018) Thus, this kind of data was not considered in our analyses (see Smith et al. 2020 and Tan et al. 2015). Briefly, in all these studies, cognitive performance was assessed using associative learning paradigms testing the ability of bees to associate an olfactory or/and a visual stimulus with an appetitive or aversive reinforcement (Giurfa, 2007). Olfactory learning was tested in 9 out of the 24 studies. These studies used olfactory learning protocols with appetitive conditioning of the extension of the proboscis of bees (PER; 17 studies) or aversive conditioning of the sting extension (SER; 2 studies). Either response was conditioned by presenting bees a conditioned stimulus (an odour) paired with an unconditioned stimulus (a reward of sucrose solution or an electric shock), for 3-15 trials in appetitive assays and 5-6 trials in aversive assays. Trainings included absolute learning (the odour is reinforced) and differential learning (an odour is reinforced, the other is not). Visual learning was tested in 5 out of 24 the studies. These studies used visual learning protocols with appetitive conditioning in a Y-maze (1 study) or on artificial flowers (1 study), or aversive conditioning with electric shocks (1 study). One of these studies applied a multimodal appetitive conditioning combining both odour and colour cues to be learnt by bees in an array of artificial flowers (Muth et al. 2019). Here again bees were tested for differential learning. The last study included a test of social recognition when placed with a conspecific (Shepherd et al. 2019).

#### Stressors

The stressor type covered different pesticides, parasites, predator odours, alarm pheromones, and heavy metal pollutants. Studies performed with pesticides whose median lethal dose (LD50; i.e. dose that kills 50% of the population) could not be identified in the literature were excluded from our final selection.

#### Exposure duration

In all these studies, stressors were applied before the cognitive tests, except in two studies in which it was used as the CS during conditioning (i.e. petrol exhaust (Leonard et al. 2019), alarm and predator pheromones (Wang et al. 2016)). We categorised the duration of exposure using the common dichotomy between acute and chronic treatments. An acute treatment was characterized by a single administration of the pesticide to each individual bee. When bees were exposed to the pesticide more than once, either as a substance present in their environment or as a food directly offered to each individual, the exposure type was considered chronic.

#### Bee genus

The bee species studied in the selected publications were the honey bees *Apis cerana* and *Apis mellifera*, and the bumblebees *Bombus impatiens* and *Bombus terrestris*. These species were not selected purposefully, but rather emerged from the refinement obtained with other inclusion criteria. All but three raw datasets were available online with the published material. Those three datasets were kindly provided by their authors. The list of the 24 selected studies is available in Table 1. The raw data are provided in Dataset S1.

### Dataset organisation and normalisation of variables

The raw data were downloaded and saved as .csv files. A new dataset (Dataset S1) was created, which combined information on the species, the cognitive task studied, the type of stressor, the type of exposure (acute/chronic), and, in the case of pesticide studies, the dose (μg/bee) or concentration (ppb). Within each study, data were grouped in different categories according to homogeneous experimental methodologies (i.e. 38 categories).

To allow comparison across studies, a z-score was calculated for each individual on its cognitive performance by applying the function ‘scale’ in R (package {base}) which uses the mean and the standard deviation of the sample to scale each element. Within each study, the function ‘scale’ was applied on the cognitive performance of bees belonging to the same category of bee species, cognitive task, stressor type and exposure type. When learning performance was measured as a binary response (e.g. success vs. failure) across multiple trials, the raw data was first used to calculate a learning score for each individual corresponding to the number of successful trials. Such a calculation was required because the variance in binary variables can be mathematically predicted from the mean and sample size and does not reflect biological variance (Supplementary Fig. S1). For pesticide studies, the dose (acute exposure) and concentration (chronic exposure) were normalized using the LD50.

Individual z-scores were used to calculate the mean and the variance of the z-scores for each control and stressed group. We thereafter refer to these variables as the “mean” and the “variance” of the cognitive performance. Each study may contain multiple control and stressed groups depending on the number of experiments performed and the number of stressors used. The final sample sizes are therefore larger than the number of studies and are displayed on the figures.

### Data analyses

All analyses were conducted in R Studio v.1.2.5033 (RStudio Team 2015). Linear mixed-effects models (LMMs; package {lme4}; Bates et al. 2015) were used to investigate the impact of stressors on the mean and the variance of the cognitive performance. The group (control vs. stressed), the type of stressor, the species or the type of tasks were defined as independent variables. The experiment’s identifier was set as random factor.

Similar models were used to assess the impact of pesticides on the mean and variance of the cognitive performance. In the subset of pesticide studies (15 studies), Pearson correlation tests were also performed to assess the relationship between the mean and the variance of the cognitive performance within control and stressed groups. LMMs were conducted to study the influence of the pesticide dose (log-transformed) on individual z-scores, with the experiment’s identifier set as random factor.

## Results

### All stressors reduced the cognitive mean but not the variance

We first explored the overall effects of stress on the cognitive mean and variance of bees across the 24 studies. As expected from previous studies, the mean cognitive performance was severely impacted by exposure to stressors (Fig. 1A). Overall, stressed bees exhibited a significantly lower mean cognitive performance than control bees (LMMs; *group effect*: F_1,90_ > 15, *P* < 0.001 for all models) irrespective of the type of stressor they were exposed to (*group*stressor effect*: F_4,90_ = 0.92, *P* = 0.454; Fig. 1A), the bee genus (*group*genus effect*: F_1,94_ = 1.23, *P* = 0.271; Fig. 1C) and the type of cognitive task (*group*task effect*: F_3,93_ = 0.84, *P* = 0.477; Fig. 1E).

**Figure 1.**
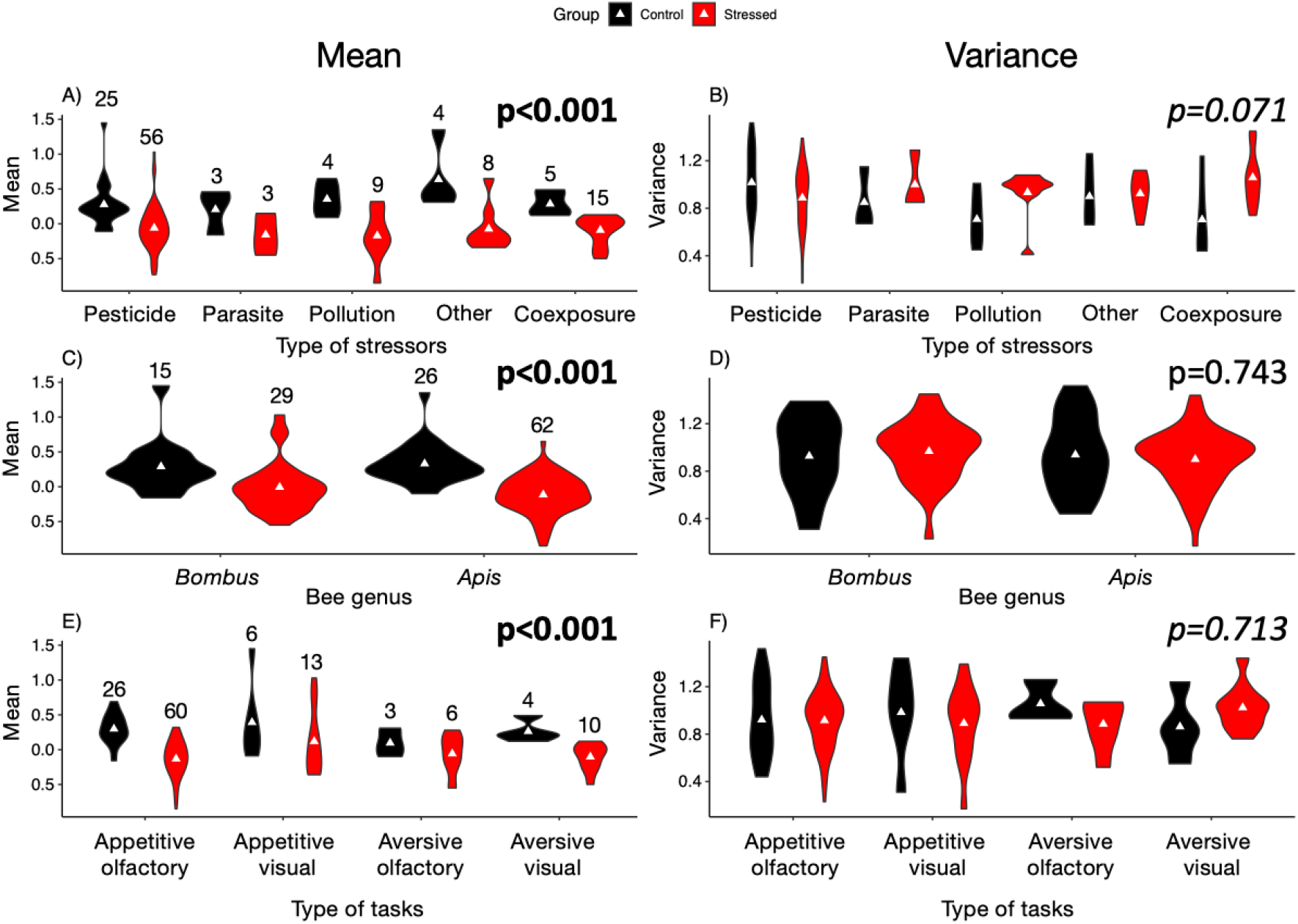
Stressors decrease the mean cognitive performance of bees, but not the variance. Violin plots showing the mean (left) and the variance (right) of the cognitive performance for control (black) and stressed (red) bees are displayed according to: **A-B)** the type of stressors; **C-D)** the bee genus; **E-F)** the type of cognitive tasks. White triangles represent the mean. Sample sizes are displayed above the violins. P-values from LMM are displayed for group effect only and are in bold when significant.

The effects of stressors on cognitive variance were less pronounced and more heterogeneous (Fig. 1B). Variance did not differ significantly between control and stressed bees (LMMs; *group effect*: F_4,122_ < 4.12, *P* > 0.05 for all models). We found no effect of the bee genus (*group*genus effect*: F_1,128_ = 0.65, *P* = 0.421; Fig. 1D) nor of the type of cognitive task (*group*task effect*: F_3,120_ = 0.75, *P* = 0.533; Fig. 1F). There was a significant interaction between exposure to stressor and the type of stressor, indicating a heterogeneous effect of stressors on the variance of the cognitive performance (*group*stressor effect*: F_4,122_ = 3.44, *P* = 0.011; Fig. 1B). While variance decreased in stressed bees exposed to pesticides, it tended to increase in stressed bees exposed to other stressor types, compared to their respective control group. Thus exposure to stressors globally reduced the cognitive performances of bees, with mixed effects on variance depending on stressor type.

### Chronic exposure to pesticides reduced cognitive mean and variance

To investigate whether stressor intensity plays a role in the differential effects of stressors observed on the variance of the cognitive performance of bees, we focused our analyses on the 15 pesticide studies of our dataset (Table 1). Pesticide studies were the most abundant in the literature and present the advantage that a normalization of stressor intensity across drugs was possible using LD50s (amount of substance necessary to kill 50% of individuals in the population) and durations of exposure (acute or chronic).

Both acute and chronic treatments reduced the mean cognitive performance of bees (Fig. 2; LMM; *acute*: F_1,34_ = 5.89, estimates±standard error: −0.232±0.095, *P* = 0.021; *chronic*: F_1,20_ = 28.69, −0.465±0.083, *P* < 0.001). They also tended to reduce the cognitive variance within populations, although to different extent. Cognitive variance of stressed bees was significantly lower than that of control bees in the chronic treatments (Fig. 2B; F_1,20_ = 10.34, estimates: −0.317±0.107, *P* < 0.01) but not in the acute treatments (Fig. 2A; F_1,47_ = 0.40, estimates: −0.005±0.078, *P* = 0.532). However, we found a positive correlation between the mean cognitive performance and its variance in both stressed groups (*acute*: r = 0.437, *P* < 0.01; *chronic*: r = 0.657, *P* < 0.005), but not in control groups (*acute*: r = 0.057, *P* = 0.868; *chronic*: r = 0.072, *P* = 0.833). This shows pesticides tended to reduce both mean and variance in the two treatments, but this effect was more pronounced for chronic exposure.

**Figure 2.**
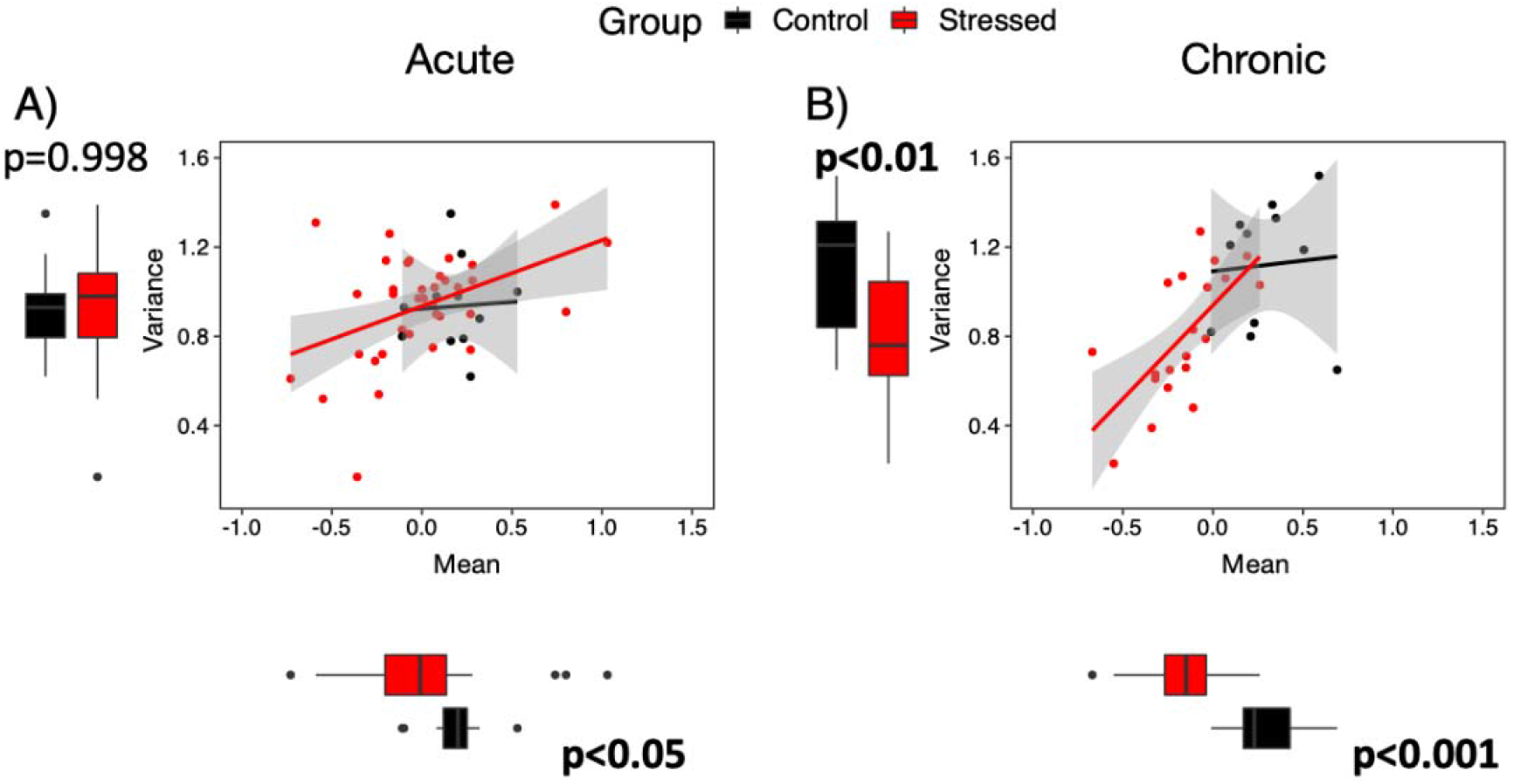
Pesticide exposure duration affects the variance of the cognitive performance. The mean and the variance of the cognitive performance are plotted for control (black) and stressed (red) bees following an **A)** acute (N = 13 controls, N = 36 stressed) or **B)** chronic (N = 11 controls, N = 20 stressed) exposure to pesticides. Horizontal and vertical boxplots represent the mean cognitive performance and its variance, respectively. P-values from LMMs are displayed for group effect only and are in bold when significant.

### High pesticide doses reduced cognitive mean and variance

To further explore whether the effect on mean and variance differed with stress magnitude, we analysed different doses and durations of pesticide exposure. A dose-dependent effect on cognitive performance was found for both acute and chronic exposure (Fig. 3). Cognitive performances (Individual z-scores) significantly decreased with increasing doses of exposure (LMM; Fig. 3A, *acute*: estimates = −0.144±0.018, *P* < 0.001, Fig. 3B; *chronic*: estimates = −0.121±0.020, p < 0.001). Interestingly, both mean and variance decreased with increasing pesticide doses for acute and chronic exposures (Figs 3C-D). This means most bees in the population tested seemed to show a decreased cognitive performance following a treatment with high pesticide doses, irrespective of exposure duration.

**Figure 3.**
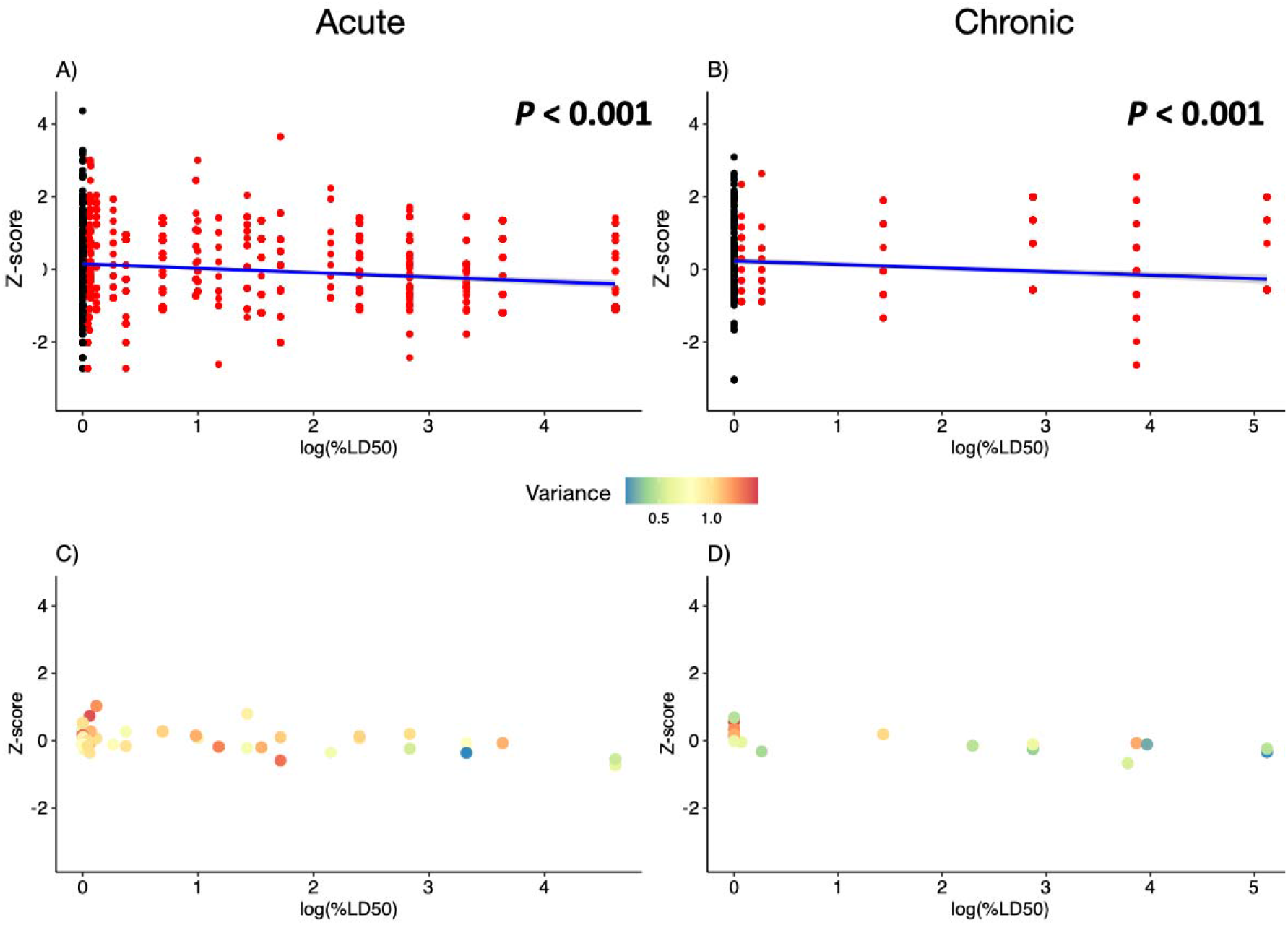
Effect of the pesticide dose on cognitive performance. Individual z-scores are plotted relative to the normalized pesticide dose (logarithm of %LD50) for **A)** acute exposure (N = 2,141 bees) and **B)** chronic exposure (N = 1,026 bees). Estimate trends are displayed in solid blue lines. Plots showing the mean cognitive performance relative to the normalized pesticide dose (logarithm of %LD50) and coloured according to variance for **C)** acute (N = 13 controls, N = 36 stressed) and **D)** chronic exposure (N = 11 controls, N = 11 stressed).

## Discussion

Many environmental stressors affect the behaviour and cognition of animals (Killen et al. 2013; Klein et al. 2017; Siviter et al. 2018; Siviter et al. 2021). While studies reporting such sublethal effects have typically focused on mean phenotypic traits, all individuals in a population are not similarly affected by stressors, and the resulting phenotypic variance may be critical for stress resilience. Here we tested this hypothesis by analysing raw datasets of 24 bee studies. We showed different effects on the cognitive mean and variance of insects exposed to stressors, depending on stress level and exposure mode, thus validating the importance of examining variance in addition to mean phenotypic traits in ecotoxicological studies.

Focusing on pesticide revealed the mean cognitive performance of bees was altered by both chronic and acute exposures. This result is consistent with a previous meta-analysis (Siviter *et al*., 2018). However, the variance in cognitive performance of bees was only decreased after a chronic exposure. This means some bees were able to better cope than others with short pesticide exposure, but not to repeated stress. This is, to our knowledge, the first study showing a differential effect of acute and chronic exposures to a stressor on learning performance in an animal. Such variance in response to stress might be due to homeostatic physiological processes that can counteract the effect of an acute exposure to the drug, which is only present in the body for a short duration (Cohen, 2006). Indeed most pesticides act on the nervous system of bees whose plasticity to maintain homeostasis is well-known (Turrigiano and Nelson, 2000; Cabirol and Haase, 2019). For instance, neurons can compensate a change in the balance between brain excitation and inhibition by modulating the efficacy of specific synapses(Pozo and Goda, 2010). As neonicotinoids activate the excitatory cholinergic neurotransmission pathway, one might expect the brain to counteract this increased excitation (Cabirol and Haase, 2019). However, the lasting presence of toxic compounds in the bodies during a chronic exposure seems to complicate the process of resilience to this stressor for most individuals.

Interestingly, for both acute and chronic pesticide exposure, the mean cognitive performance and its variance decreased with increasing doses of toxic compounds. The positive correlation between the mean and the variance is consistent with this finding: the more a stressor affects the mean, the more it affects the variance. This advocates for the use of low pesticide concentrations in the field. Reducing use to doses having sublethal effects on pest insects would still protect crops when pest density is low and thereby would be less damaging to non-target insects (Colin, Monchanin, *et al*., 2020).

Altogether, our results thus suggest that an acute exposure to low pesticide doses is the least damaging for bee populations. Indeed, despite the reduced mean cognitive performance, an unaltered variance of the learning performance following pesticide exposure means that some individuals may have maintained sufficient cognitive abilities to support efficient foraging (Klein *et al*., 2017). Cognitive and behavioural variance is thought to be particularly important for populations resilience after environmental changes (Jandt *et al*., 2014) as it augments the probability that some individuals display adapted behaviour to the new environmental conditions. In group-living species, such as social insects, the high diversity of behavioural phenotypes within colonies is known to improve the efficiency of collective decision-making and the ability of groups to find optimal solutions to changing conditions (Burns and Dyer, 2008; Michelena *et al*., 2010).

In nature, bees often encounter pesticides over long time periods especially when colonies are located near treated crops and in the hive due to the residues present in the honey and wax (Godfray *et al*., 2014, 2015; Tsvetkov *et al*., 2017). The consequences of such a chronic exposure to pesticides are often not a priority in risk assessment procedures. Policy regulations in the European Union and in the US regarding the commercialization of new plant protection products (PPPs) ask for acute toxicity assays on bees and other non-target animals before asking for chronic toxicity assays (EPPO, 1992, 2010; U.S. Environmental Protection Agency and Code of Federal Regulations (CFR), 2010). Only when acute toxicity is significant would a chronic toxicity assay be performed. Although the European Food Safety Authority recommends the inclusion of chronic exposure assays earlier in the risk assessment procedure, such assays are not yet mandatory (EFSA, 2013). The effects of PPPs that will be encountered chronically in the field might therefore be underestimated. Note that the fact similar results were obtained in *Bombus* and *Apis* confirms honey bees are overall suitable surrogates for non-*Apis* species in regulatory risk assessments of pesticide toxicity (Arena and Sgolastra, 2014; Heard *et al*., 2017; Thompson and Pamminger, 2019), as currently considered by the European commission (EPPO 2010). This is true at least when exploring general trends. But these results must then be complemented on non-*Apis* bee species that may vary in their sensitivity to pesticides (Arena and Sgolastra, 2014).

Overall, our study revealed a differential effect of chronic and acute exposures to pesticides as well as an important influence of the stressor intensity on the proportion of individuals that might be impacted. Focusing on variance helped identify acute stress conditions bees may be able to cope with, which could not be done by looking at the mean only. Interestingly all types of stressors did not similarly influence bee cognition. While the mean was severely impacted by all stressors, variance seemed to increase in some non-pesticide stressors. This positive effect could be triggered by the relatively small sample sizes found for some stressors (N ≤ 5 for the control groups used to assess the effect of parasites, pollution, and co-exposures). But if it is confirmed, this means stress can favour the diversification of cognitive abilities (Badyaev, 2005), an observation already made in rodents where low intensity stressors can have beneficial effects on the cognitive performance (Hurtubise and Howland, 2016). These intriguing effects of stress on cognitive traits demonstrate the importance of considering phenotypic variance in future analyses of the impact of environmental stressors on animals. We hope such approach can be generalised to assess more thoroughly the hazardous nature of the stressors and identify the modes of exposure that might be less damaging for wild populations. Future investigations should also consider the possible interaction between agrochemicals, which have synergistic effects on bee mortality, but antagonistic effects on behaviour when looking at the mean only (Siviter *et al*., 2021). Ultimately the results of such studies should lead to explicit guidelines for farmers on the safe use of these toxic substances.

## Authors’ contribution

AC, CP and ML designed the study. AC, TGM and CM collected the data. TGM processed the data and prepared dataset. CM, TGM and CP analysed the data. AC wrote the first draft of the manuscript. All authors substantially contributed to revisions.

## Acknowledgements

We thank Jacques Gautrais for discussions about mean and variance correlations, and Dara Stanley and Ken Tan for providing raw data of their studies.

## Data availability statement

Raw data are available in Dataset S1 (.xlsx file). The data supporting the results will be archived in Dryad Digital Repository upon publication of the manuscript.

## Supplementary materials

**Supplementary figure S1.**
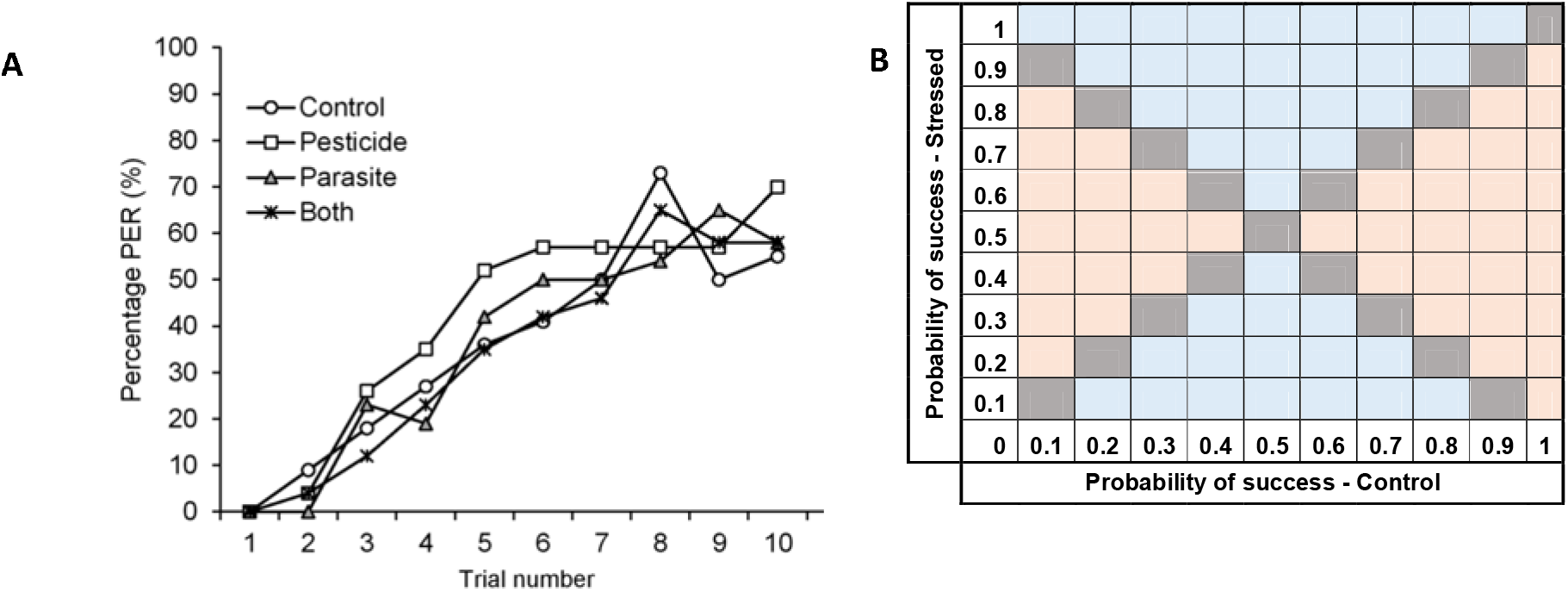
Analysis of variance in studies with binary data. **(A)** Impact of a pesticide and parasite on bumblebees’ learning performance measured with a classical conditioning of the proboscis extension response (PER) (from Piiroinen et al. 2016). The percentage of individuals that extended the proboscis in response to the conditioned stimulus (i.e percentage of learners) is plotted across 10 successive learning trials. **(B)** Matrix representing the impact of a stressor on the variance in learning performance. For an equal sample size in the control and treatment groups, the impact of the treatment on variance can be calculated using the mean of each group. An increased (orange) or decreased variance (blue) can be predicted.

